# Fiber alignment alone is insufficient to predict cellular contact guidance responses in engineered 3D collagen hydrogels

**DOI:** 10.64898/2026.09.29.755189

**Authors:** Poorya Esmaili, Tresa M. Elias, Indranil M. Joshi, Richard A. Simon, Dinindu De Silva, Sami Farajollahi, Olivia M. Harris, Edward B. Brown, Vinay V. Abhyankar

## Abstract

In breast cancer, collagen fiber alignment and the second harmonic generation forward-to-backward ratio (F/B), an optical measurement sensitive to fibril-scale organization *within* collagen fibers, have each been associated with metastatic outcome. Although both are measured at the tumor-stromal interface, *in vitro* studies have largely examined them separately, because experimental systems have not allowed the two to be varied independently. To address this technical gap, we used a microfluidic biofabrication approach to generate paired aligned and unaligned regions within 3D collagen hydrogels via extensional strain and varied measured F/B across gels by changing the pre-gel pH. The resulting library covered alignment values reported in breast tumors and F/B values associated with poorer metastatic outcome in invasive ductal carcinoma, at matched matrix stiffness, pore diameter, and fiber fraction. In MDA-MB-231 cells, alignment biased migration along the fiber axis without changing migration speed, as expected, but the magnitude of the contact guidance response varied with measured F/B. In contrast, MCF-7 cells showed little overall contact guidance, yet migrated ∼30% faster in high F/B than low F/B gels in both aligned and unaligned regions. Together, these results show that cells migrated differently in matrices with similar fiber alignment. Thus, including measured F/B may provide additional information to help anticipate how cells migrate in collagen environments that appear similar based on alignment alone.

## 1. Introduction

In their native tissue environment, cells are surrounded by the extracellular matrix (ECM), a complex biopolymer network that provides mechanical support and instructive cues, influencing a wide range of cell behaviors ^1–3^. Type I collagen is the most abundant structural protein in the ECM, and its organization is established through a hierarchical self-assembly process. Collagen molecules associate into fibrils tens of nanometers in diameter, which pack within a semiregular lattice to form fibers one to two micrometers across. These fibers then crosslink to form dynamic networks that span millimeters to centimeters ^4–7^. Thus, the assembly process presents distinct features at different length scales. The degree of fiber alignment describes the organization of the collagen network, while the size, arrangement, and packing of fibrils within a fiber define its internal structure. Both features can be assessed using standard optical imaging techniques ^7–9^.

At the network scale, fiber alignment is a well-established migration cue. As shown in **Figure 1**, during tumor invasion, cancer cells migrate through the collagen-rich stroma toward vascular and lymphatic vessels, with aligned fibers biasing migration along the fiber axis ^10–14^. Aligned fibers oriented perpendicular to the tumor–stromal boundary are prognostic of metastatic outcome in breast and pancreatic tumors, and overall fiber alignment in the tumor bulk is predictive and prognostic as well ^15–21^. Similar contact guidance responses contribute to endothelial and immune cell migration during angiogenesis and immune trafficking ^22–24^ and have been reproduced in vitro using micropatterned grooves, electro-spun fiber arrays, and aligned 3D collagen hydrogels ^25–29^. At the level of the individual fiber, the ratio of forward-to backward propagating second harmonic generation signal (F/B) provides an optical measure sensitive to fibril diameter, spacing, and packing order ^5,7,30–32^. Similar to alignment, measured F/B values have been associated with patient outcome across multiple tumor types, with lower F/B at the tumor–stromal interface associated with shorter metastasis-free survival in invasive ductal carcinoma (IDC) ^33–36^. The same measurement in the tumor bulk is not prognostic _34_.

**Figure 1.**
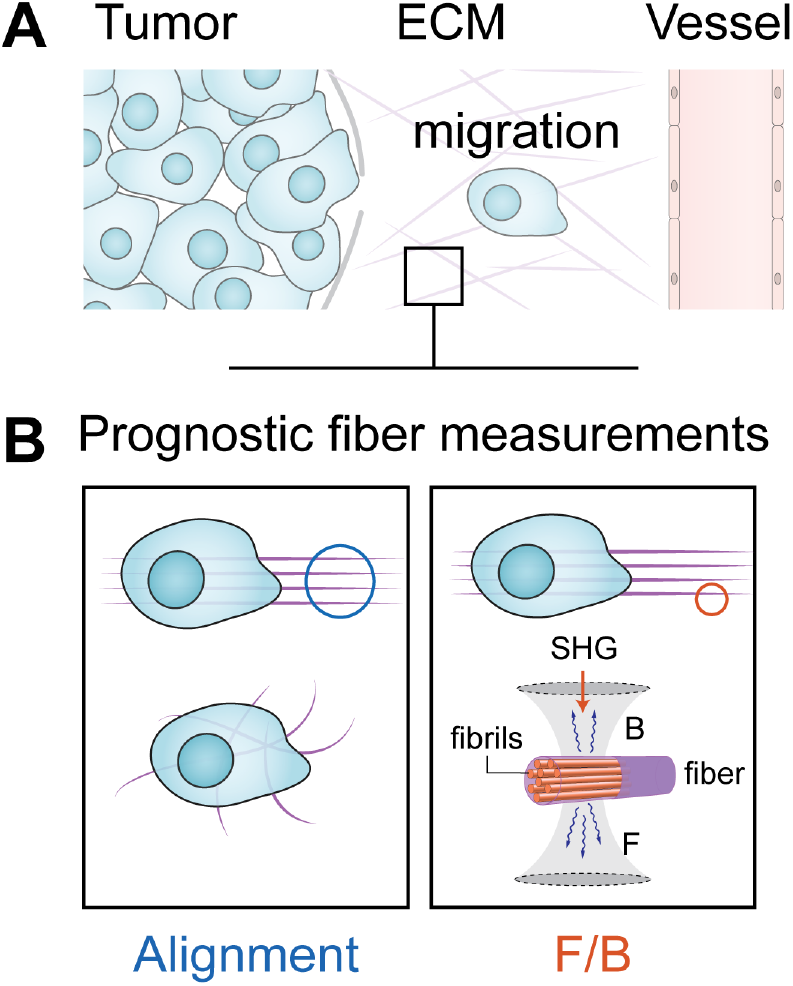
(A) Schematic of cancer cell migration from a tumor into the surrounding collagen-rich stroma. (B) Fiber alignment describes organization of the collagen network, whereas the second harmonic generation forward-to-backward ratio (F/B) is an optical measurement sensitive to fibril-scale organization within collagen fibers. Both measurements have been associated with breast cancer outcome.

Despite their clinical relevance, fiber alignment and measured F/B have largely been examined separately in vitro. Cell migration responses across gels with different F/B values have been studied without controlling fiber alignment ^37,38^, while aligned 3D collagen hydrogels have been used to study contact guidance without measuring F/B ^12,28,29^. Previous experimental systems have not allowed fiber alignment and measured F/B to be varied independently, preventing direct comparison of migration responses across matrices with similar fiber alignment but different measured F/B, or similar measured F/B but different alignment.

To address this gap, we used a microfluidic platform to engineer a library of 3D collagen hydrogels containing both aligned and unaligned regions within individual gels and different measured F/B values across gels. The library spanned fiber alignment and F/B values observed in breast tumor tissue ^34,35,39^. We then tested whether fiber alignment alone was sufficient to anticipate migration behavior across this library and whether measured F/B provided additional context for two cell populations, MDA-MB-231 and MCF-7, with distinct migratory phenotypes.

## 2. Materials and Methods

### 2.1 Experimental design

Each collagen gel was fabricated in a segmented microfluidic channel that produced aligned and unaligned fiber regions within the same device, and each gel had a single measured F/B value (**Figure 2A**). Fiber alignment therefore varied within a gel and F/B varied between gels. Pre-gel pH was adjusted before injection to generate gels spanning different measured F/B values ^37,38^, with lower pH corresponding to lower measured F/B.

**Figure 2.**
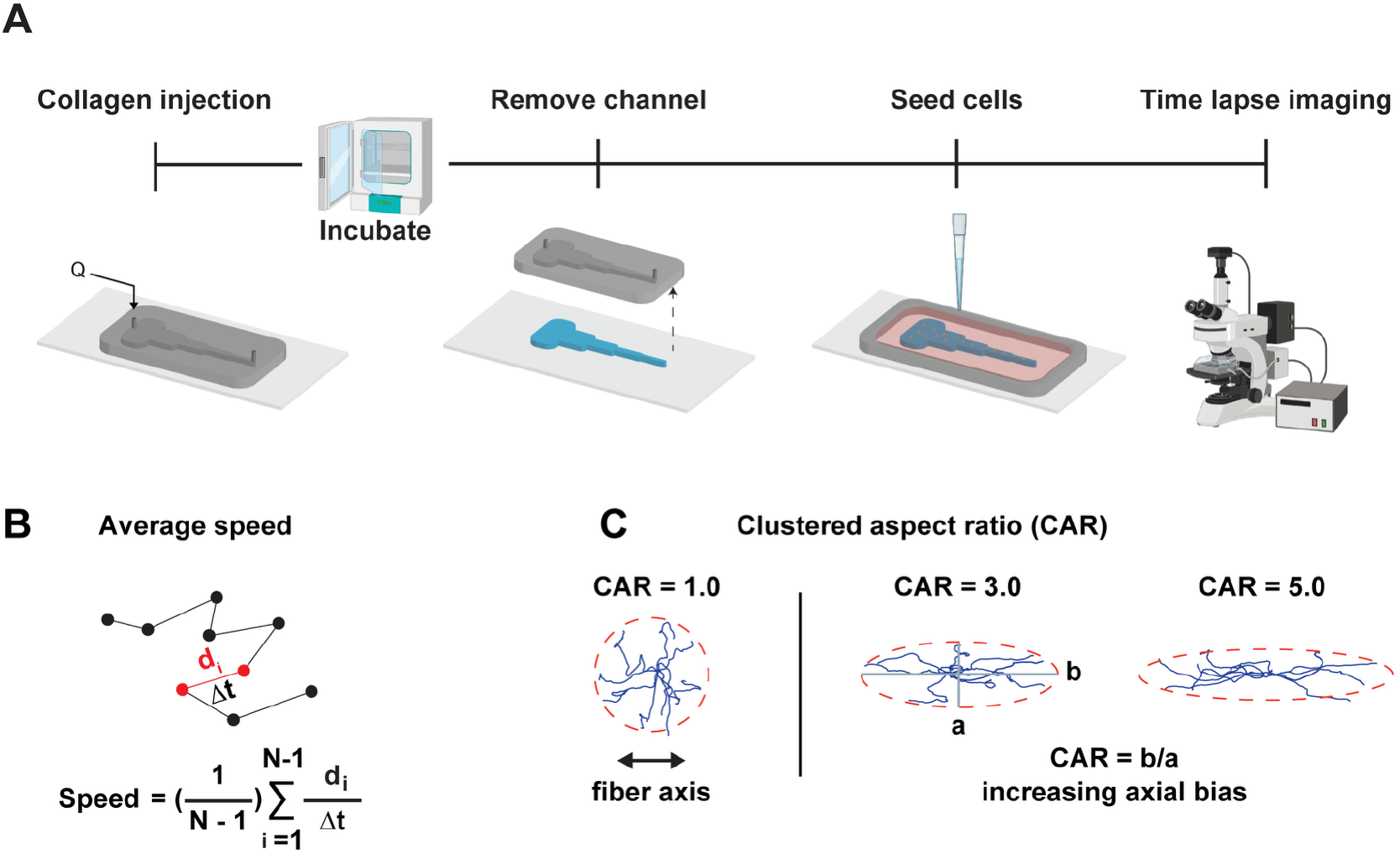
Experimental workflow and migration metrics. (A) Collagen was injected into a PDMS microfluidic channel and polymerized for 1 h at room temperature followed by 1 h at 37 °C. The PDMS channel was then removed to expose the 3D collagen gel. An acrylic border was placed around the gel to retain culture medium, cells were seeded onto the collagen, and time-lapse imaging was performed for 6 h at 2-min intervals. (B) Migration speed is the mean frame-to-frame displacement, d_i_, divided by the imaging interval Δt. (C) Clustered Aspect Ratio (CAR) quantifies axial bias in the trajectory distribution. Cell trajectories are translated to a common origin and fitted with a 99% confidence ellipse. CAR is the ratio of the major to minor ellipse axes, with values closer to 1 indicating weaker axial bias and larger values indicating stronger axial bias. Example trajectory distributions illustrate increasing CAR values.

Two to three imaging windows were collected from each region of a gel, with approximately 25 tracked cells per window. Window values were averaged to give one aligned and one unaligned value per gel for each migration measurement. The gel was therefore the experimental unit for all migration analyses. The final datasets comprised 38 gels for MDA-MB-231 cells and 37 for MCF-7 cells, for 68 and 67 gel-level alignment regions, respectively. Thirty gels in each dataset contributed both an aligned and an unaligned region. The remaining eight MDA-MB-231 gels and seven MCF-7 gels contributed a single alignment class because damage to the complementary region prevented cell seeding. These damaged regions were excluded from migration analysis, but the gels were retained. Gels contributing both regions were used for the aligned–unaligned comparisons, and all gels were included in the comparisons among F/B categories.

### 2.2 Microfluidic device fabrication

The microfluidic guidance-channel mold was fabricated using standard soft lithography ^40–42^. SU-8 3050 negative photoresist (Kayaku Advanced Materials, MA, USA) was spin-coated onto a 100-mm silicon wafer (University Wafers, MA, USA) to a target thickness of 200 µm, defining the microchannel height. The photoresist was soft-baked at 95 °C for 60 min, exposed to 365-nm ultraviolet light (250 mJ/cm^2^) through a high-resolution photomask, post-baked at 95 °C for 5 min, developed in SU-8 developer, rinsed with isopropanol, and dried with pressurized air. PDMS channels were fabricated from Sylgard 184 base and crosslinker (Dow Corning, MI, USA) mixed at 10:1, degassed, poured over the mold, and cured at 80 °C for 1 h. After cooling, the channels were removed and cleaned with 70% ethanol, and inlet and outlet ports were created using 0.8-mm biopsy punches.

Glass coverslips were functionalized with poly(octadecene-alt-maleic anhydride) (POMA; Millipore Sigma, MA, USA) to improve collagen adhesion ^41,43,44^. Coverslips were sonicated in isopropyl alcohol, rinsed with deionized water, dried, heated at 100 °C, and activated using oxygen plasma at 600 mTorr for 1 min (Harrick Plasma, NY, USA). Activated coverslips were immersed in 2% v/v aminopropyltriethoxysilane (APTES; Millipore Sigma, MA, USA) in acetone, rinsed, dried, spin-coated with POMA dissolved in tetrahydrofuran (Millipore Sigma, MA, USA), and cured at 120 °C for 1 h.

### 2.3 Collagen preparation and pre-gel pH

Bovine telo Type I collagen (Advanced BioMatrix, CA, USA) was prepared at a final concentration of 2.7 mg/mL on ice by mixing stock collagen with 10× PBS, deionized water, and 0.1 M NaOH. The volumes of NaOH and water were adjusted to produce pre-gel pH values from 7.0 to 10.7, confirmed using a calibrated pH probe (Orion Star A211, Thermo Fisher Scientific, USA).

### 2.4 Hydrogel formation and generation of fiber alignment

Fiber alignment was generated using a segmented microfluidic channel as previously described ^40,41^. The channel comprised five sequential 5 mm long segments of uniform height (200 µm) with stepwise reductions in width (10, 5, 2.5, 1.25, and 0.75 mm). The change in geometry between segments produced localized extensional strain during injection, generating segments of different alignment within a single gel. Observation regions (665 µm × 665 µm) were positioned centrally within each segment.

Before injection, PDMS channels were passivated in 4% bovine serum albumin in PBS for 2 h at 4 °C, rinsed, and dried. Collagen was loaded into a 1 mL syringe fitted with a 20-gauge dispensing needle and injected at 250 µL min^-1^ using a syringe pump (New Era Pump Systems, NY, USA). Devices were incubated for 1 h at room temperature and 1 h at 37 °C, after which the PDMS channel was removed to expose the collagen surface. We have previously shown that collagen alignment is unaffected by channel removal ^40,41^.

Following channel removal, an acrylic border was added as a media reservoir, gels were equilibrated at 37 °C and 5% CO_2_ in culture media before cell seeding. Therefore, all gels were at physiological pH at the time of seeding, regardless of the pre-gel pH used during fabrication.

### 2.5 Measurement of fiber alignment and F/B

Following the migration experiments, gels were fixed with 4% paraformaldehyde (Thermo Fisher Scientific, MA, USA) and imaged by confocal reflectance microscopy on a Leica SP5 laser-scanning confocal microscope using a 40× water-immersion objective. Collagen fibers were imaged in reflectance mode using a 488-nm laser. Z-stacks of 13 slices were acquired 10–20 µm below the gel surface, corresponding to the depth at which cells were tracked, and projected using average-intensity projection in Fiji ^45^. Fiber angles were extracted using CT-FIRE ^9^, binned in 15° increments, and the coefficient of alignment (CoA) was calculated as the fraction of fibers oriented within ±15° of the modal angle. At CoA ≥ 0.5, most fibers in a region fall within a single 30° band; below 0.5, no single direction accounts for the majority of fibers. Regions were categorized as unaligned (CoA < 0.5) or aligned (CoA ≥ 0.5) ^39^.

SHG imaging was performed using 810-nm excitation with circular polarization. Backward-propagating SHG was collected through the excitation objective and forward-propagating SHG through a condenser, using matched 405/30 bandpass filters and photomultiplier tubes. Optical settings and collection geometry were identical for all gels. Forward and backward images were thresholded independently using an adaptive thresholding algorithm and combined to define a collagen mask, using the same thresholding method as the clinical studies against which these values are compared ^34,35^. The mean forward-to-backward intensity ratio within the mask was reported as the gel F/B value ^33,34^.

Gels were assigned to low (2 ≤ F/B < 6), medium (6 ≤ F/B < 10), or high (10 ≤ F/B < 15) categories. Alignment was treated as two classes because collagen fibers oriented perpendicular to the tumor–stromal boundary are scored as present or absent in clinical practice, where their presence is prognostic of survival ^15,19,21^. Measured F/B is reported as a continuous value ^34^ and was stratified into three categories within the range.

Spatial uniformity of F/B was assessed in a separate set of seven gels. F/B was measured in five regions spanning the width of the segmented channel and normalized to the mean F/B of each gel. F/B did not differ among regions (**Figure S1**), and each gel in the library was assigned a single measured F/B value.

### 2.6 Characterization of matrix properties

Stiffness was measured using a Piuma nanoindenter (Optics11, Amsterdam, Netherlands) with a 27 µm radius spherical probe and a 0.016-N m^-1^ cantilever. Sixteen positions were sampled per gel, and force-displacement curves were fitted to a Hertz contact model. Pore diameter and fiber fraction were quantified from confocal reflectance images using established methods. After contrast correction and binarization, pore area was calculated from non-fiber regions and fiber fraction as the ratio of fiber-positive pixels to total pixels ^46^. Six gels were measured per condition, and five in the low-F/B group.

### 2.7 Cell culture

MDA-MB-231 cells (GenTarget, San Diego, CA, USA) were expanded for three passages after thawing and used for fewer than eight total passages. Cells were maintained in DMEM (Gibco, Thermo Fisher Scientific, MA, USA; Cat. No. 11320033) supplemented with 10% fetal bovine serum (Thermo Fisher Scientific, USA; Cat. No. A5670701) and 1% penicillin/streptomycin (Gibco, Thermo Fisher Scientific, MA, USA; Cat. No. 15140122).

MCF-7 cells (GenTarget, San Diego, CA, USA) were expanded for three passages after thawing and used for fewer than eight total passages. Cells were maintained in MEM (Gibco, Thermo Fisher Scientific, MA, USA; Cat. No. 11095080) supplemented with 1 mM non-essential amino acids (Gibco, Thermo Fisher Scientific, MA, USA; Cat. No. 11140050), sodium pyruvate (Gibco, Thermo Fisher Scientific, MA, USA; Cat. No. 11360070), 10% fetal bovine serum, and 1% penicillin/streptomycin. All cells were maintained at 37 °C and 5% CO_2_.

### 2.8 Migration assay and migration measurements

Cells were labeled with SPY650-DNA cell tracker dye (1:1000; Cytoskeleton Inc., Denver, CO, USA) and incubated for 1 h at 37 °C, then seeded onto the equilibrated gels at 10,000 cells/cm^2^. Cells infiltrated 15 to 20 µm into the collagen network and were tracked within the matrix.

Time-lapse imaging was performed on an Olympus IX-81 inverted microscope using a 20× objective at 37 °C and 5% CO_2_ in a stage top incubator (Okolab, CA, USA). Samples equilibrated on the stage for 1 h before acquisition. Images were collected for 6 h at 2 min intervals, stabilized in Fiji, and trajectories generated using TrackMate ^47^. Trajectory coordinates were analyzed using custom scripts in MATLAB R2026a (MathWorks, Natick, MA, USA).

Migration speed was calculated as the mean frame-to-frame displacement divided by the frame interval (**Figure 2B**). Values were averaged across tracked cells within an imaging window and then across windows of the same alignment class (aligned or unaligned), giving one aligned and one unaligned speed value per gel.

To quantify the tendency of cells to move along the fiber direction, trajectories within each imaging window were translated to a common origin and a covariance-based 99% confidence ellipse was fitted to the centered distribution. The Clustered Aspect Ratio (CAR) was calculated as the ratio of the major (b) to minor (a) ellipse axes and averaged across windows of the same alignment class in each gel (**Figure 2C**). CAR values closer to 1 indicate weaker axial bias, and larger values indicate stronger bias along the fiber direction.

### 2.9 Statistical analysis

Because aligned and unaligned regions were generated within the same gel, alignment was compared within gels and F/B between gels. Data were analyzed in GraphPad Prism 9 using a two-factor mixed-effects model fitted by restricted maximum likelihood, testing alignment class, F/B category, and their interaction, with values matched by gel. The alignment term tested the overall difference between aligned and unaligned regions, the F/B term tested differences among F/B categories, and the alignment by F/B interaction tested whether the aligned–unaligned difference varied among F/B categories. This fit was used in place of a repeated-measures two-way ANOVA, which would have excluded those 15 gels that contributed a single alignment class. Group values are reported as model-estimated means with 95% confidence intervals, which account for the matching and can differ slightly from the simple average of the plotted points. Tukey-adjusted pairwise comparisons identified differences among specific matrix combinations, with p < 0.05 considered statistically significant. Full model results and adjusted comparisons are provided in **Tables S2** and **S3**.

Normality was assessed using the Shapiro-Wilk test. Two-group comparisons used parametric or non-parametric tests according to distributional assumptions. Multi-group comparisons of gel structural and mechanical properties used ordinary one-way ANOVA with Tukey’s post hoc test when equal variances were assumed, Brown-Forsythe and Welch ANOVA with Dunnett’s T3 post hoc test when variances were unequal, or the Kruskal–Wallis test with Dunn’s post hoc test when normality was not met.

## 3. Results

### 3.1 Engineered hydrogel library spans the alignment and F/B ranges measured in breast tumors

To explore migration responses to combinations of prognostic collagen matrix properties, we began by establishing a library of 3D hydrogels in which fiber alignment varied within gels and measured F/B varied across gels. Alignment was controlled using our established extensional strain-based microfluidic platform ^40,41^, which created defined unaligned (CoA < 0.5) and aligned (CoA ≥ 0.5) regions within the same channel. Pre-gel pH was varied to generate gels spanning different measured F/B values ^37,38^. As shown in **Figure S1**, measured F/B did not differ across measurement regions within the device, and each gel was therefore assigned a single measured F/B value. The composition of the engineered gel library is shown in **Figure 3**. CoA values spanned 0.25 to 0.68, covering the range measured in human breast tumor specimens (0.34 to 0.67) ^39^, while F/B values spanned 2.9 to 14.7, overlapping the range reported at the tumor– stromal interface associated with poorer metastatic outcome in IDC ^34,35^. Gels were grouped into low (2 ≤ F/B < 6), medium (6 ≤ F/B < 10), and high (10 ≤ F/B < 15) categories. The mean CoA in unaligned and aligned regions ranged from 0.33 to 0.38 and 0.56 to 0.62, respectively, across all F/B categories. The library therefore contained aligned and unaligned regions at every F/B level.

**Figure 3.**
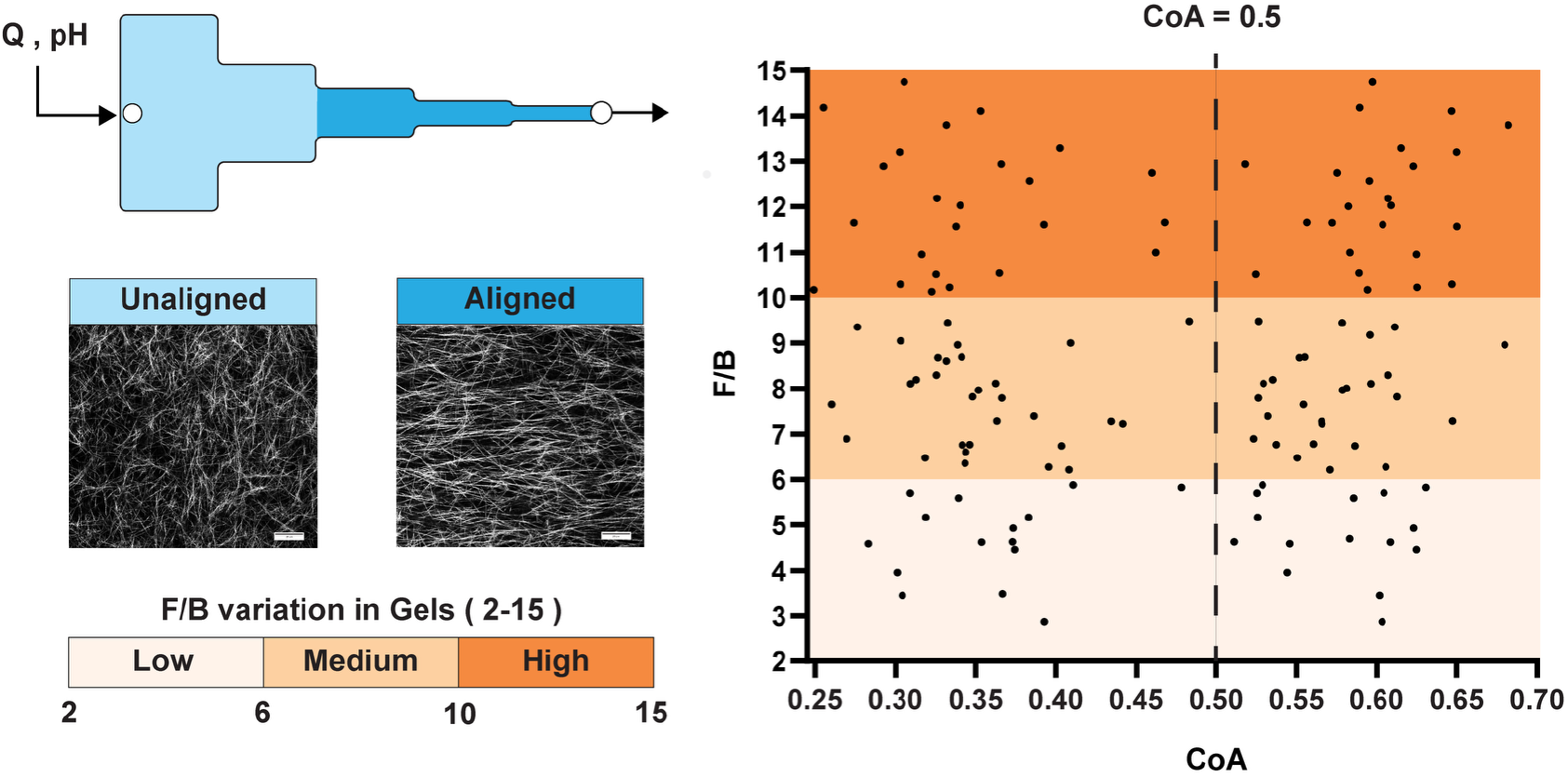
The alignment-F/B experimental space. The segmented microfluidic channel generated regions of different fiber alignment within each gel, while pre-gel pH was varied across gels to generate different measured F/B values. Measured CoA and F/B values across the hydrogel library spanned CoA from 0.25 to 0.68 and F/B from 2.9 to 14.7. Unaligned (CoA < 0.5) and aligned (CoA ≥ 0.5) regions were represented across low (2 ≤ F/B < 6), medium (6 ≤ F/B < 10), and high (10 ≤ F/B < 15) F/B gels. Representative confocal reflectance images show unaligned and aligned collagen. Fiber alignment is visible by reflectance microscopy, whereas measured F/B is obtained by SHG and is sensitive to fibril-scale organization within collagen fibers.

Because the two cancer cell populations in this study were investigated in independently fabricated gels, we next tested whether the gels themselves differed in ways that could confound our interpretation of the migration results. In gels used for the MDA-MB-231 studies, CoA was less than 0.48 in unaligned regions and was at least 0.52 in aligned regions, with a mean within-gel Δ_CoA_ = 0.24 ± 0.07 across the 30 gels containing both regions. The corresponding values in the MCF-7 gels were less than 0.48 and at least 0.51, with a mean within-gel Δ_CoA_ = 0.23 ± 0.07 across 30 gels. Measured F/B ranged from 3.5 to 14.2 in the MCF-7 gels, matching the gels used in the MDA-MB-231 experiments, and all three F/B categories were represented in both studies (**Table S1**). The two gel sets therefore covered similar alignment and F/B ranges.

We also tested whether other matrix properties differed across alignment class and F/B categories. The collagen gels had a stiffness of approximately 110 Pa, a mean pore diameter of 2.6 µm, and a fiber fraction of 0.19, with no detectable differences between aligned and unaligned regions or across F/B categories (**Figure 4**). With the gel library established and potential confounding factors assessed, we next examined cell migration behaviors within the gels.

**Figure 4.**
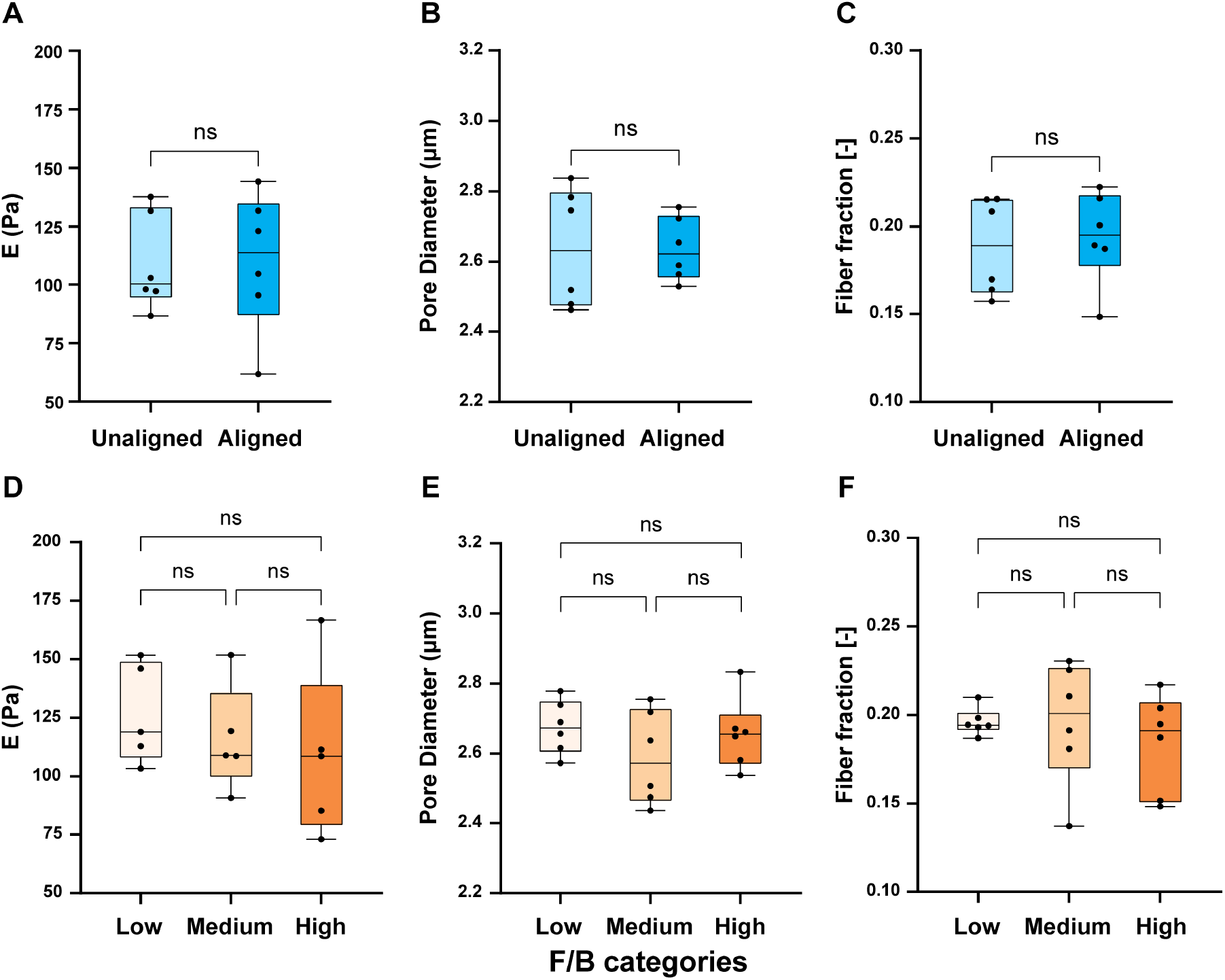
Stiffness, pore diameter, and fiber fraction across the library. Stiffness, pore diameter, and fiber fraction were compared between unaligned and aligned regions (A–C) and across low, medium, and high F/B gels (D–F). No detectable differences were observed between alignment classes or among F/B categories for any measured property. Points represent independent gels; Box plots indicate the median and interquartile range, with whiskers representing minimum and maximum values. n = 6 gels per group except D panel, for which n = 5. Statistical tests were selected according to distributional and variance assumptions as described in Section 2.9; ns, p ≥ 0.05.

### 3.2 Alignment increases axial bias in MDA-MB-231 migration, and the magnitude of the response varies with measured F/B

We first examined MDA-MB-231 cells because their contact guidance response to aligned features is well characterized ^11–13^. We quantified migration speed and CAR (**Figure 2B**,**C**) to determine whether the magnitude of this response differed across measured F/B categories.

As shown in **Figure 5A**, MDA-MB-231 mean migration speed ranged from 0.38 to 0.40 µm min^-1^ across alignment and F/B combinations, with no difference between paired aligned and unaligned regions (p = 0.97) or among F/B categories (p = 0.76). The aligned-unaligned difference in speed was also similar across F/B categories (alignment by F/B interaction, p = 0.85). Because contact guidance is bidirectional, CAR was measured from an ellipse fitted to the track distribution rather than from net displacement. In aligned regions, the major axis of the ellipse coincided with the fiber axis. CAR was 1.68 in unaligned regions and 2.61 in aligned regions, with an aligned– unaligned difference of 0.94 CAR units (95% CI, 0.70 to 1.17; p < 0.0001) (see **Figure 5B**). Alignment therefore increased axial bias in MDA-MB-231 migration without affecting speed ^12^.

**Figure 5.**
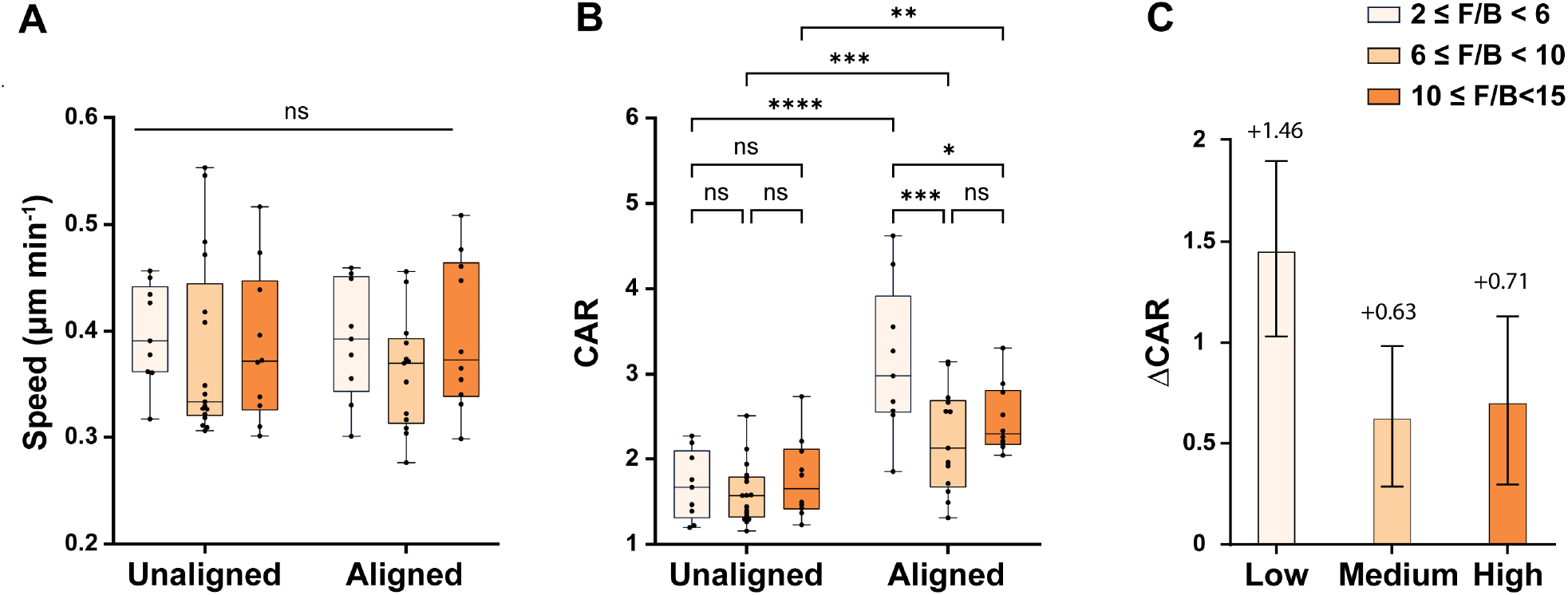
MDA-MB-231 migration behavior across the hydrogel library. (A) Migration speed and (B) CAR in unaligned (CoA < 0.5) and aligned (CoA ≥ 0.5) regions grouped by measured F/B category. F/B categories were low (2 ≤ F/B < 6), medium (6 ≤ F/B < 10), and high (10 ≤ F/B < 15). Points represent gel-level measurements, boxes indicate the median and interquartile range, and whiskers indicate the observed range. Migration speed did not differ between alignment classes or among F/B categories. Alignment increased CAR, but the magnitude of the increase differed across measured F/B categories.(C) Aligned–unaligned CAR difference (ΔCAR) within each F/B category estimated from the two-factor mixed-effects model. Bars show model-estimated differences with 95% confidence intervals. ΔCAR was greatest in low F/B gels, with smaller responses in medium and high F/B gels (alignment by F/B interaction, p = 0.012). n = 38 gels. Pairwise comparisons were Tukey-adjusted; ns, p ≥ 0.05; *p < 0.05; **p < 0.01; ***p < 0.001; ****p < 0.0001.

Alignment increased CAR in every F/B category, but the magnitude of that increase differed across measured F/B categories (alignment by F/B interaction, p = 0.012). ΔCAR was 1.46 in low F/B gels, larger than in either medium or high F/B gels (0.63 and 0.71, respectively), which did not differ from each other (**Figure 5C**). Within unaligned regions, CAR did not differ among F/B categories. The variation in ΔCAR across F/B categories was not explained by differences in the degree of fiber alignment. Aligned regions in low and medium F/B gels reached comparable CoA values (0.57 ± 0.04 and 0.56 ± 0.03) but differed more than twofold in ΔCAR. High F/B gels contained the most aligned regions (CoA 0.62 ± 0.04) but produced a ΔCAR of 0.71, approximately 50% lower than the ΔCAR of 1.46 in low F/B gels. Thus, similar or greater fiber alignment did not produce equivalent contact guidance responses.

### 3.3 MCF-7 migration speed varies with measured F/B rather than with alignment

We next examined MCF-7 cells, which retain a more epithelial and less invasive migratory phenotype than MDA-MB-231 cells ^27,48,49^, providing a contrasting system to study whether the relationships between alignment, measured F/B, and migration were consistent across cell populations with different invasive properties.

MCF-7 cells exhibited no meaningful contact guidance response. CAR ranged from 1.29 to 1.43 across all alignment classes and F/B combinations (**Figure 6A**), and the overall aligned-unaligned difference was −0.005 CAR units (p = 0.87). An alignment by F/B interaction was detected (p = 0.027), indicating that the aligned–unaligned CAR difference varied among F/B categories. However, the category-specific differences were small and did not represent a consistent increase in CAR with alignment. The aligned−unaligned difference within high F/B gels (0.10 CAR units, p = 0.041) and the difference between medium and high F/B within unaligned regions (0.14 CAR units, p = 0.035) were statistically significant, but neither exceeded 0.15 CAR units, compared to the 0.94 CAR unit in the alignment response for MDA-MB-231 cells (**Table S3**). Alignment, therefore, did not produce a meaningful axial bias in this cell population.

**Figure 6.**
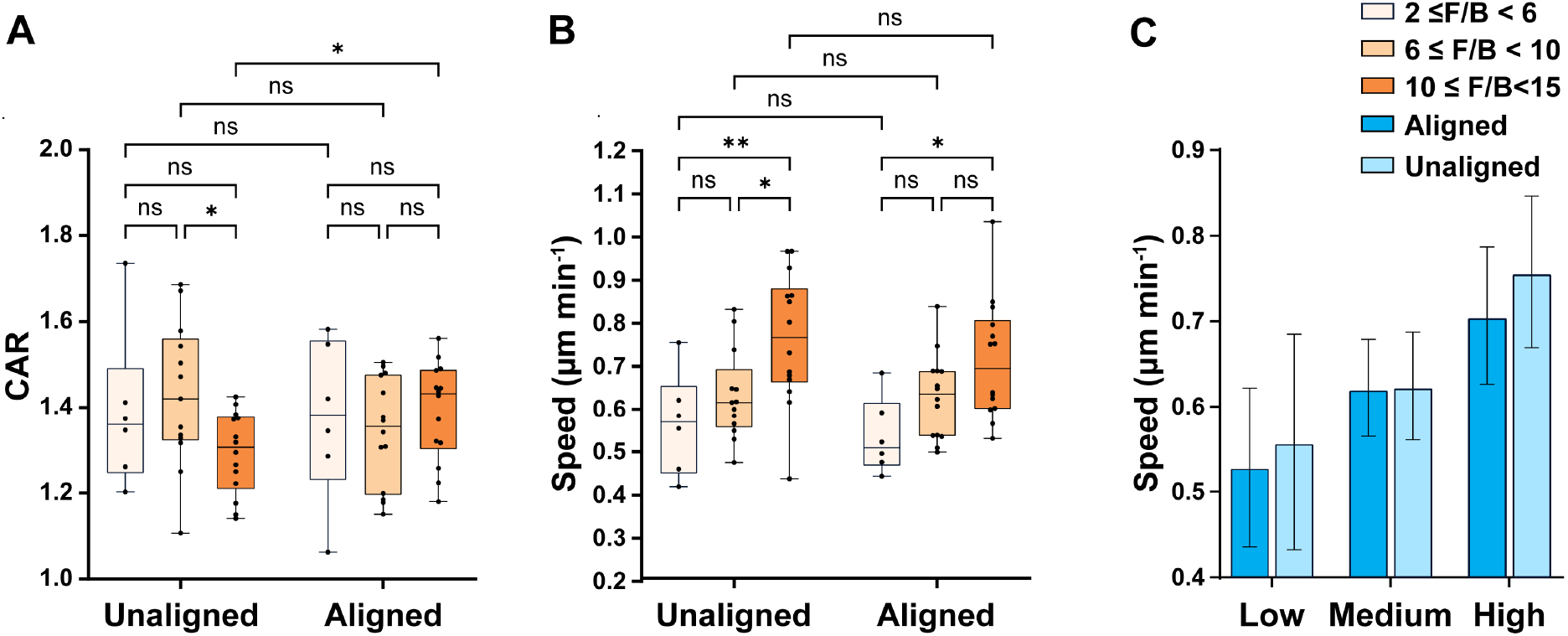
Migration of MCF-7 cells across the hydrogel library. (A) CAR and (B) speed in unaligned (CoA < 0.5) and aligned (CoA ≥ 0.5) regions grouped by measured F/B category. F/B categories were low (2 ≤ F/B < 6), medium (6 ≤ F/B < 10), and high (10 ≤ F/B < 15). Points represent gel-level measurements; boxes indicate the median and interquartile range and whiskers indicate the observed range. CAR remained within a narrow range. Although the aligned–unaligned CAR difference varied among F/B categories (alignment by F/B interaction, p = 0.027), the category-specific differences were small, and alignment did not produce a consistent increase in CAR. (C) Model-estimated migration speed across F/B categories for aligned and unaligned regions; error bars indicate 95% confidence intervals. Migration speed differed among F/B categories (p = 0.002), with higher speed in high F/B than low F/B gels in both aligned and unaligned regions. The pattern across F/B categories was similar in the two alignment classes (alignment by F/B interaction, p = 0.53). n = 37 gels. Pairwise comparisons were Tukey-adjusted; ns, p ≥ 0.05; *p < 0.05; **p < 0.01; ***p < 0.001; ****p < 0.0001.

As shown in **Figure 6B**, MCF-7 migration speed differed among measured F/B categories (p = 0.002). In unaligned regions, speed was 0.56 µm min^−1^ in low F/B gels, 0.62 µm min^−1^ in medium F/B gels, and 0.76 µm min^−1^ in high F/B gels. Speed was 0.19 µm min^−1^ higher in high F/B than low F/B gels (p = 0.004), corresponding to a 34% increase. In aligned regions, speed increased from 0.55 µm min^−1^ in low F/B gels to 0.71 µm min^−1^ in high F/B gels, a difference of 0.16 µm min^−1^ (p = 0.023), corresponding to a 29% increase (**Figure 6C)**.

Speed did not differ between aligned and unaligned regions (p = 0.33), and the pattern across F/B categories was similar in both alignment classes (alignment by F/B interaction, p = 0.53). Thus, differences in MCF-7 speed among F/B categories were present whether the collagen was aligned or unaligned. Together, measured F/B provided additional information about the magnitude of alignment-associated axial bias in MDA-MB-231 cells and migration speed in MCF-7 cells.

## 4. Discussion

Fiber alignment is widely used as a single parameter to predict migration responses to collagen architecture in tumor tissue and engineered matrices ^12,15–17,28,29^. In our system, however, fiber alignment alone was not sufficient to predict migration behavior across collagen matrices with different measured F/B values. In MDA-MB-231 cells, alignment strongly increased CAR, but the magnitude of this increase differed across measured F/B categories while migration speed remained nearly unchanged. In MCF-7 cells, alignment produced little change in CAR, whereas migration speed was approximately 30% higher in high F/B than low F/B gels in both aligned and unaligned regions. The weak contact guidance response of MCF-7 cells is consistent with prior studies showing relatively limited guidance of epithelial breast cancer cells in aligned fiber matrices ^27^. Higher F/B has previously been associated with greater motility of mouse 4T1 cells in collagen gels without controlled alignment ^37,38^. To our knowledge, this relationship has not been examined in human breast cancer cells or with controlled alignment.

The difference between the two cell lines may reflect their position on the epithelial–mesenchymal spectrum. MDA-MB-231 cells are mesenchymal, whereas MCF-7 cells retain a more epithelial phenotype characterized by high E-cadherin and cytokeratin expression ^50–52^. EMT has been reported to increase susceptibility to contact guidance in cells moving on aligned collagen-coated PCL fibers ^27^. Guidance along aligned structures has also been associated with anisotropic focal adhesion organization ^25^ and linked to vimentin-dependent focal adhesion maturation ^53^. Whether the differences observed here are associated with EMT state could be tested using our biofabrication approach with a single cell population driven into epithelial, intermediate, and mesenchymal states.

Stiffness, pore diameter, and fiber fraction were matched across the library, making these measured matrix properties unlikely to account for the migration differences observed. Measured F/B is an optical measurement sensitive to fibril-scale organization within collagen fibers and provided information not captured by alignment, stiffness, pore diameter, or fiber fraction. Because F/B is sensitive to multiple aspects of fibril-scale organization, it should not be interpreted as a direct cellular cue, but rather as an additional measured metric to consider when comparing collagen environments.

Two factors limit the scope of the current findings, and both can be addressed within our platform. The gels had a stiffness of approximately 110 Pa, which is well below the stiffness reported for normal and malignant breast tissue ^54,55^. Because the channel is removed to expose the gel, external crosslinking can be used to increase stiffness without changing how the fibers were formed, allowing the relationships identified in this work to be tested across physiological and pathophysiological stiffness ranges. Our findings also come from two established breast cancer cell lines, and extending the comparison to additional lines and to primary tumor cells would establish how broadly F/B relates to different migration behaviors.

For MDA-MB-231 cells, the magnitude of the alignment-associated increase in CAR differed across measured F/B categories, with the strongest contact guidance response occurring in low F/B gels. These gels occupied the portion of the measured F/B range associated with poorer metastatic outcome in IDC ^34,35^. In our system, measured F/B was associated with the magnitude of the guidance response only where fibers were aligned. Clinically, F/B is prognostic at the tumor–stromal interface, where aligned fibers are present, and not in the tumor bulk ^15,21,34^. Our results do not establish a mechanism linking these clinical observations to cell migration, but they show that cells exhibited different contact guidance responses in matrices with similar or greater fiber alignment, within a clinically relevant F/B range. Because fiber alignment and F/B can both be measured from SHG images of the same tissue region ^35,56^, future studies can directly test whether considering both measurements improves the association with metastatic outcome compared with either measurement alone.

## 5. Conclusions

We engineered a library of 3D collagen hydrogels in which fiber alignment varied within gels and measured F/B varied between gels, spanning values relevant to breast tumor tissue. This approach enables migration to be compared across gels that differ in fiber alignment and measured F/B while retaining similar measured stiffness, pore diameter, and fiber fraction. Alignment strongly increased CAR in MDA-MB-231 cells, but the magnitude of this response differed across measured F/B categories. MCF-7 cells showed little overall contact guidance, yet migration speed differed across F/B categories in both aligned and unaligned regions. Together, these results show that fiber alignment alone does not fully describe migration behavior across collagen environments with different measured F/B values. Considering measured F/B alongside fiber alignment therefore provides additional information for characterizing migration responses in engineered collagen matrices.

## Acknowledgments

The authors thank Dr. Hyla Sweet for assistance with confocal microscopy and members of the Biological Microsystems Laboratory for insightful discussions, as well as Showmick Paul for assistance with SHG imaging. This work was supported in part by the National Science Foundation through grant numbers 2150798 (VVA) and 2150799 (EBB). The opinions, findings, conclusions, or recommendations expressed are our own views and do not necessarily reflect the views of the National Science Foundation.

## Supplementary Information

**Table S1.**
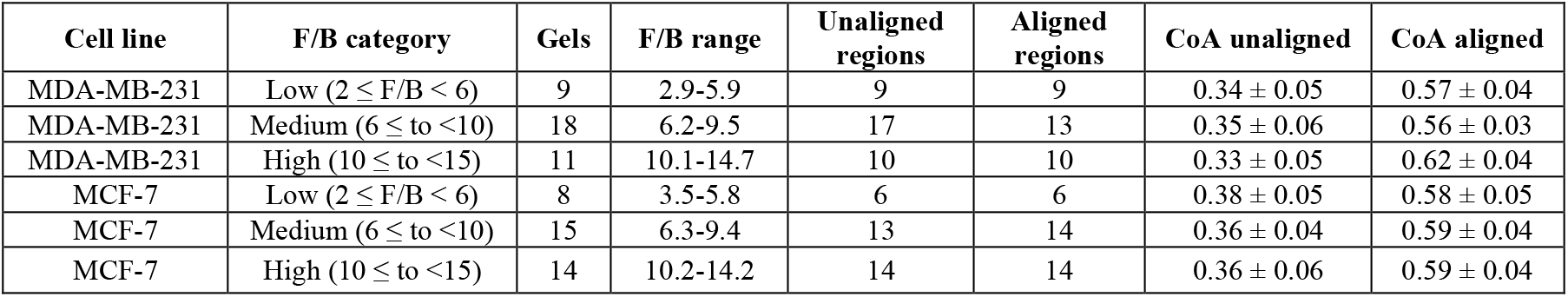
Distribution of gels and alignment regions across measured F/B categories. Gels were assigned to F/B categories after measurement. Gels contributing only one alignment class were retained in the mixed-effects model. CoA values are mean ± SD.

**Table S2.**
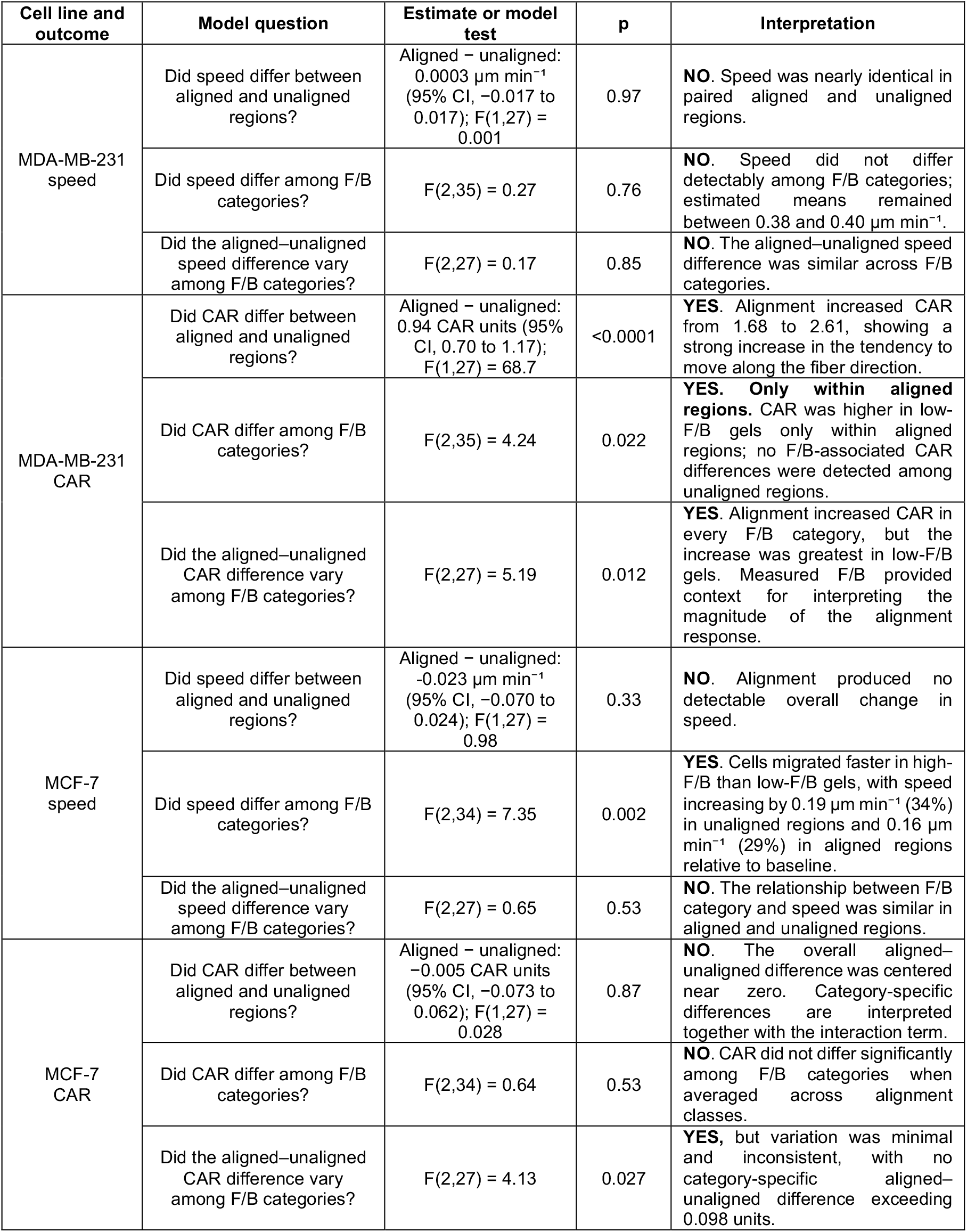
Mixed-effects model results for migration speed and CAR. The table relates each model question to a numerical result and its interpretation. Alignment class, F/B category, and their interaction were tested with values matched by gel. Estimated means and differences were calculated from full-precision model output and rounded independently. Displayed differences may therefore not equal subtraction of the rounded means.

**Table S3.**
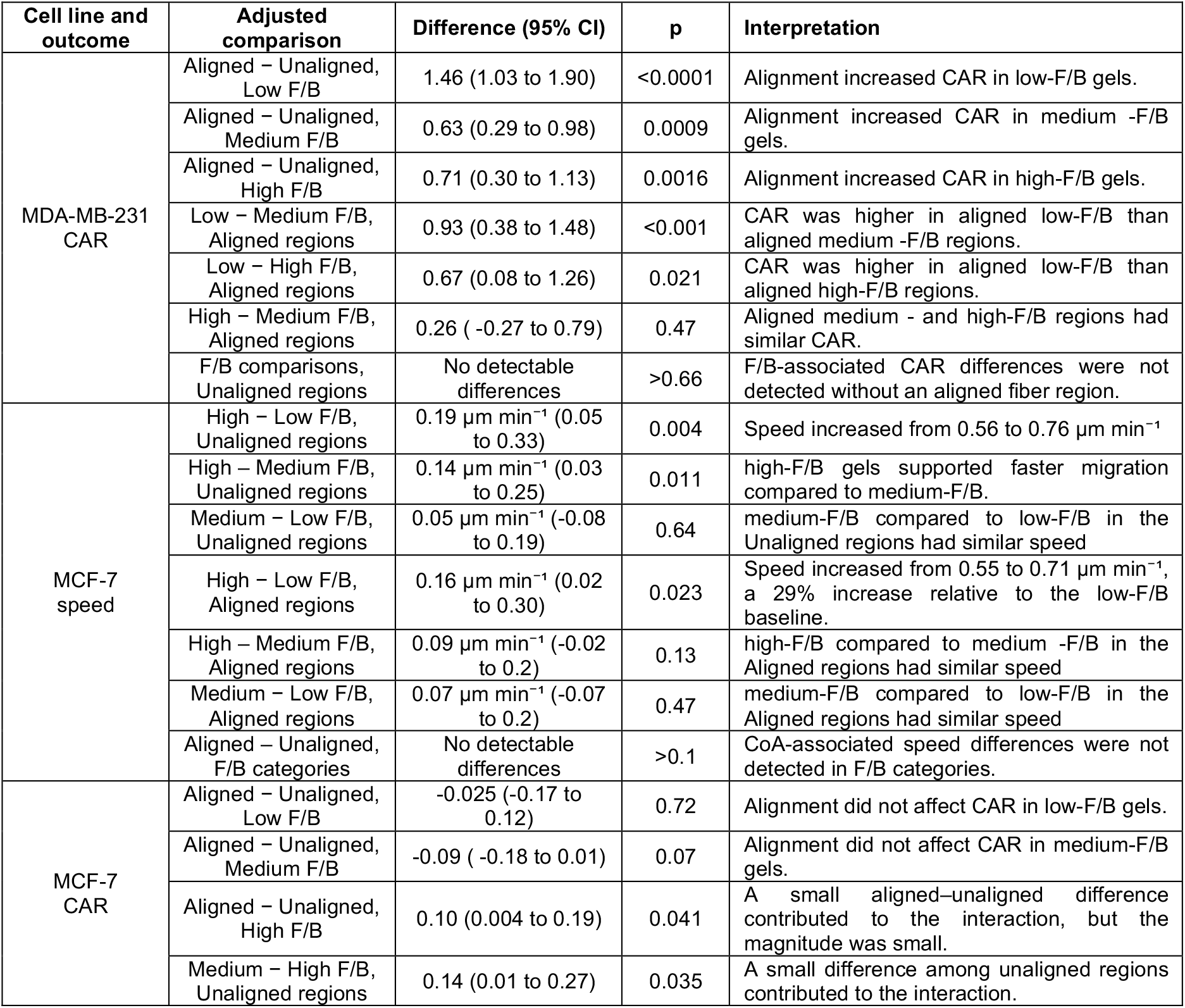
Pairwise comparisons supporting the main-text interpretation. Differences, 95% confidence intervals, and p-values are Tukey-adjusted for multiple comparisons.

| Cell line and outcome | Adjusted comparison | Difference (95% CI) | p | Interpretation |
| --- | --- | --- | --- | --- |
| MDA-MB-231 CAR | Aligned – Unaligned, Low F/B | 1.46 (1.03 to 1.90) | <0.0001 | Alignment increased CAR in low-F/B gels. |
|  | Aligned – Unaligned, Medium F/B | 0.63 (0.29 to 0.98) | 0.0009 | Alignment increased CAR in medium -F/B gels. |
|  | Aligned – Unaligned, High F/B | 0.71 (0.30 to 1.13) | 0.0016 | Alignment increased CAR in high-F/B gels. |
|  | Low – Medium F/B, Aligned regions | 0.93 (0.38 to 1.48) | <0.001 | CAR was higher in aligned low-F/B than aligned medium -F/B regions. |
|  | Low – High F/B, Aligned regions | 0.67 (0.08 to 1.26) | 0.021 | CAR was higher in aligned low-F/B than aligned high-F/B regions. |
|  | High – Medium F/B, Aligned regions | 0.26 ( -0.27 to 0.79) | 0.47 | Aligned medium - and high-F/B regions had similar CAR. |
|  | F/B comparisons, Unaligned regions | No detectable differences | >0.66 | F/B-associated CAR differences were not detected without an aligned fiber region. |
| MCF-7 speed | High – Low F/B, Unaligned regions | 0.19 $\mu\text{m min}^{-1}$ (0.05 to 0.33) | 0.004 | Speed increased from 0.56 to 0.76 $\mu\text{m min}^{-1}$ |
| | High – Medium F/B, Unaligned regions | 0.14 $\mu\text{m min}^{-1}$ (0.03 to 0.25) | 0.011 | high-F/B gels supported faster migration compared to medium-F/B. |
| | Medium – Low F/B, Unaligned regions | 0.05 $\mu\text{m min}^{-1}$ (-0.08 to 0.19) | 0.64 | medium-F/B compared to low-F/B in the Unaligned regions had similar speed |
| | High – Low F/B, Aligned regions | 0.16 $\mu\text{m min}^{-1}$ (0.02 to 0.30) | 0.023 | Speed increased from 0.55 to 0.71 $\mu\text{m min}^{-1}$, a 29% increase relative to the low-F/B baseline. |
| | High – Medium F/B, Aligned regions | 0.09 $\mu\text{m min}^{-1}$ (-0.02 to 0.2) | 0.13 | high-F/B compared to medium -F/B in the Aligned regions had similar speed |
| | Medium – Low F/B, Aligned regions | 0.07 $\mu\text{m min}^{-1}$ (-0.07 to 0.2) | 0.47 | medium-F/B compared to low-F/B in the Aligned regions had similar speed |
|  | Aligned – Unaligned, F/B categories | No detectable differences | >0.1 | CoA-associated speed differences were not detected in F/B categories. |
| MCF-7 CAR | Aligned – Unaligned, Low F/B | -0.025 (-0.17 to 0.12) | 0.72 | Alignment did not affect CAR in low-F/B gels. |
|  | Aligned – Unaligned, Medium F/B | -0.09 ( -0.18 to 0.01) | 0.07 | Alignment did not affect CAR in medium-F/B gels. |
|  | Aligned – Unaligned, High F/B | 0.10 (0.004 to 0.19) | 0.041 | A small aligned–unaligned difference contributed to the interaction, but the magnitude was small. |
|  | Medium – High F/B, Unaligned regions | 0.14 (0.01 to 0.27) | 0.035 | A small difference among unaligned regions contributed to the interaction. |

**Figure S1.**
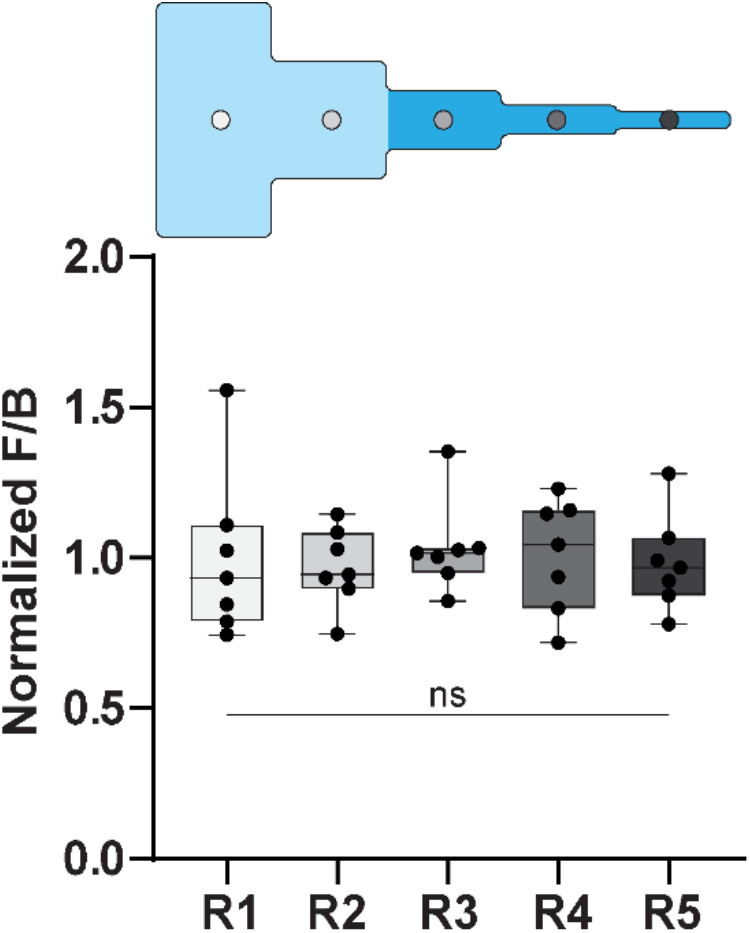
Spatial uniformity of measured F/B within collagen gels. F/B was measured at five positions spanning the segmented channel and normalized to the mean F/B of each gel. Each point represents the normalized measurement from one gel at the indicated position; boxes indicate the median and interquartile range and whiskers indicate the observed range. No detectable positional differences in normalized F/B were observed (n=7, one-way ANOVA, p>0.05, ns = not significant).

## References

1. Spill F, Reynolds DS, Kamm RD, Zaman MH. Impact of the physical microenvironment on tumor progression and metastasis. Current Opinion in Biotechnology. 2016;40:41–48. doi:10.1016/j.copbio.2016.02.007

2. Doyle AD, Carvajal N, Jin A, Matsumoto K, Yamada KM. Local 3D matrix microenvironment regulates cell migration through spatiotemporal dynamics of contractility-dependent adhesions. Nat Commun. 2015;6(1):8720. doi:10.1038/ncomms9720

3. Carey SP, Goldblatt ZE, Martin KE, Romero B, Williams RM, Reinhart-King CA. Local extracellular matrix alignment directs cellular protrusion dynamics and migration through Rac1 and FAK. Integr Biol (Camb). 2016;8(8):821–835. doi:10.1039/c6ib00030d

4. Shoulders MD, Raines RT. Collagen Structure and Stability. Annual Review of Biochemistry. 2009;78:929–958. doi:10.1146/annurev.biochem.77.032207.120833

5. Williams RM, Zipfel WR, Webb WW. Interpreting Second-Harmonic Generation Images of Collagen I Fibrils. Biophysical Journal. 2005;88(2):1377–1386. doi:10.1529/biophysj.104.047308

6. Kadler KE, Holmes DF, Trotter JA, Chapman JA. Collagen fibril formation. Biochem J. 1996;316(1):1–11. doi:10.1042/bj3160001

7. Chen X, Nadiarynkh O, Plotnikov S, Campagnola PJ. Second harmonic generation microscopy for quantitative analysis of collagen fibrillar structure. Nat Protoc. 2012;7(4):654–669. doi:10.1038/nprot.2012.009

8. Liu Y, Keikhosravi A, Mehta GS, Drifka CR, Eliceiri KW. Methods for Quantifying Fibrillar Collagen Alignment. In: Rittié L, ed. Fibrosis: Methods and Protocols. Springer; 2017:429–451. doi:10.1007/978-1-4939-7113-8_28

9. Bredfeldt JS, Liu Y, Pehlke CA, et al. Computational segmentation of collagen fibers from second-harmonic generation images of breast cancer. J Biomed Opt. 2014;19(1):016007. doi:10.1117/1.JBO.19.1.016007

10. Provenzano PP, Eliceiri KW, Campbell JM, Inman DR, White JG, Keely PJ. Collagen reorganization at the tumor-stromal interface facilitates local invasion. BMC Medicine. 2006;4:38. doi:10.1186/1741-7015-4-38

11. Provenzano PP, Inman DR, Eliceiri KW, Trier SM, Keely PJ. Contact Guidance Mediated Three-Dimensional Cell Migration is Regulated by Rho/ROCK-Dependent Matrix Reorganization. Biophysical Journal. 2008;95(11):11. doi:10.1529/biophysj.108.133116

12. Riching KM, Cox BL, Salick MR, et al. 3D Collagen Alignment Limits Protrusions to Enhance Breast Cancer Cell Persistence. Biophysical Journal. 2014;107(11):2546–2558. doi:10.1016/j.bpj.2014.10.035

13. Ray A, Slama ZM, Morford RK, Madden SA, Provenzano PP. Enhanced Directional Migration of Cancer Stem Cells in 3D Aligned Collagen Matrices. Biophys J. 2017;112(5):1023–1036. doi:10.1016/j.bpj.2017.01.007

14. Han W, Chen S, Yuan W, et al. Oriented collagen fibers direct tumor cell intravasation. Proc Natl Acad Sci U S A. 2016;113(40):11208–11213. doi:10.1073/pnas.1610347113

15. Conklin MW, Eickhoff JC, Riching KM, et al. Aligned Collagen Is a Prognostic Signature for Survival in Human Breast Carcinoma. Am J Pathol. 2011;178(3):3. doi:10.1016/j.ajpath.2010.11.076

16. Xi G, Guo W, Kang D, et al. Large-scale tumor-associated collagen signatures identify high-risk breast cancer patients. Theranostics. 2021;11(7):3229–3243. doi:10.7150/thno.55921

17. Li H, Bera K, Toro P, et al. Collagen fiber orientation disorder from H&E images is prognostic for early stage breast cancer: clinical trial validation. npj Breast Cancer. 2021;7(1):104. doi:10.1038/s41523-021-00310-z

18. Dekker TJA, Charehbili A, Smit VTHBM, et al. Disorganised stroma determined on pre-treatment breast cancer biopsies is associated with poor response to neoadjuvant chemotherapy: Results from the NEOZOTAC trial. Molecular Oncology. 2015;9(6):1120–1128. doi:10.1016/j.molonc.2015.02.001

19. Drifka CR, Loeffler AG, Mathewson K, et al. Highly aligned stromal collagen is a negative prognostic factor following pancreatic ductal adenocarcinoma resection. Oncotarget. 2016;7(46):76197–76213. doi:10.18632/oncotarget.12772

20. Natal RA, Vassallo J, Paiva GR, et al. Collagen analysis by second-harmonic generation microscopy predicts outcome of luminal breast cancer. Tumour Biol. 2018;40(4):1010428318770953. doi:10.1177/1010428318770953

21. Esbona K, Yi Y, Saha S, et al. The Presence of Cyclooxygenase 2, Tumor-Associated Macrophages, and Collagen Alignment as Prognostic Markers for Invasive Breast Carcinoma Patients. The American Journal of Pathology. 2018;188(3):559–573. doi:10.1016/j.ajpath.2017.10.025

22. Salmon H, Franciszkiewicz K, Damotte D, et al. Matrix architecture defines the preferential localization and migration of T cells into the stroma of human lung tumors. J Clin Invest. 2012;122(3):899–910. doi:10.1172/JCI45817

23. Bauer AL, Jackson TL, Jiang Y. Topography of Extracellular Matrix Mediates Vascular Morphogenesis and Migration Speeds in Angiogenesis. PLOS Computational Biology. 2009;5(7):e1000445. doi:10.1371/journal.pcbi.1000445

24. Pruitt HC, Lewis D, Ciccaglione M, et al. Collagen fiber structure guides 3D motility of cytotoxic T lymphocytes. Matrix Biology. 2020;85-86:147–159. doi:10.1016/j.matbio.2019.02.003

25. Ray A, Lee O, Win Z, et al. Anisotropic forces from spatially constrained focal adhesions mediate contact guidance directed cell migration. Nat Commun. 2017;8(1):14923. doi:10.1038/ncomms14923

26. Sundararaghavan HG, Saunders RL, Hammer DA, Burdick JA. Fiber alignment directs cell motility over chemotactic gradients. Biotechnology and Bioengineering. 2013;110(4):1249–1254. doi:10.1002/bit.24788

27. Isert L, Passi M, Freystetter B, et al. Cellular EMT-status governs contact guidance in an electrospun TACS-mimicking in vitro model. Materials Today Bio. 2025;30:101401. doi:10.1016/j.mtbio.2024.101401

28. Fraley SI, Wu P hsun, He L, et al. Three-dimensional matrix fiber alignment modulates cell migration and MT1-MMP utility by spatially and temporally directing protrusions. Sci Rep. 2015;5(1):1. doi:10.1038/srep14580

29. Zanotelli MR, Miller JP, Wang W, et al. Tension directs cancer cell migration over fiber alignment through energy minimization. Biomaterials. 2024;311:122682. doi:10.1016/j.biomaterials.2024.122682

30. LaComb R, Nadiarnykh O, Townsend SS, Campagnola PJ. Phase matching considerations in second harmonic generation from tissues: Effects on emission directionality, conversion efficiency and observed morphology. Optics Communications. 2008;281(7):1823–1832. doi:10.1016/j.optcom.2007.10.040

31. Burke K, Brown E. The Use of Second Harmonic Generation to Image the Extracellular Matrix During Tumor Progression. IntraVital. 2014;3(3):e984509. doi:10.4161/21659087.2014.984509

32. Ajeti V, Nadiarnykh O, Ponik SM, Keely PJ, Eliceiri KW, Campagnola PJ. Structural changes in mixed Col I/Col V collagen gels probed by SHG microscopy: implications for probing stromal alterations in human breast cancer. Biomed Opt Express. 2011;2(8):2307–2316. doi:10.1364/BOE.2.002307

33. Burke K, Smid M, Dawes RP, et al. Using second harmonic generation to predict patient outcome in solid tumors. BMC Cancer. 2015;15(1):929. doi:10.1186/s12885-015-1911-8

34. Desa DE, Strawderman RL, Wu W, et al. Intratumoral heterogeneity of second-harmonic generation scattering from tumor collagen and its effects on metastatic risk prediction. BMC Cancer. 2020;20(1):1217. doi:10.1186/s12885-020-07713-4

35. Elias TM, Desa DE, Brown EB IV, et al. Exploring racial differences in second-harmonic-generation–based prognostic indicators of metastasis in breast and colon cancer. Biophotonics Discov. 2025;2(2):022703. doi:10.1117/1.BIOS.2.2.022703

36. Desa DE, Bhanote M, Hill RL, et al. Second-harmonic generation directionality is associated with neoadjuvant chemotherapy response in breast cancer core needle biopsies. J Biomed Opt. 2019;24(8):086503. doi:10.1117/1.JBO.24.8.086503

37. Burke KA, Dawes RP, Cheema MK, et al. Second-harmonic generation scattering directionality predicts tumor cell motility in collagen gels. J Biomed Opt. 2015;20(5):051024. doi:10.1117/1.JBO.20.5.051024

38. Burke KA, Dawes RP, Cheema MK, Perry S, Brown EB III. Second-harmonic generation reveals a relationship between metastatic potential and collagen fiber structure. In: Multiphoton Microscopy in the Biomedical Sciences XIV. Vol 8948. SPIE; 2014:32–36. doi:10.1117/12.2041027

39. Joshi IM, Mansouri M, Ahmed A, et al. Microengineering 3D Collagen Matrices with Tumor-Mimetic Gradients in Fiber Alignment. Advanced Functional Materials. 2024;34(13):2308071. doi:10.1002/adfm.202308071

40. Ahmed A, Mansouri M, Joshi IM, et al. Local extensional flows promote long-range fiber alignment in 3D collagen hydrogels. Biofabrication. 2022;14(3):035019. doi:10.1088/1758-5090/ac7824

41. Ahmed A, Joshi IM, Larson S, et al. Microengineered 3D Collagen Gels with Independently Tunable Fiber Anisotropy and Directionality. Advanced Materials Technologies. 2021;6(4):2001186. doi:10.1002/admt.202001186

42. Duffy DC, McDonald JC, Schueller OJA, Whitesides GM. Rapid Prototyping of Microfluidic Systems in Poly(dimethylsiloxane). Anal Chem. 1998;70(23):4974–4984. doi:10.1021/ac980656z

43. Pompe T, Zschoche S, Herold N, et al. Maleic Anhydride CopolymersA Versatile Platform for Molecular Biosurface Engineering. Biomacromolecules. 2003;4(4):1072–1079. doi:10.1021/bm034071c

44. Pompe T, Renner L, Grimmer M, Herold N, Werner C. Functional Films of Maleic Anhydride Copolymers under Physiological Conditions. Macromolecular Bioscience. 2005;5(9):890–895. doi:10.1002/mabi.200500097

45. Schindelin J, Arganda-Carreras I, Frise E, et al. Fiji: an open-source platform for biological-image analysis. Nat Methods. 2012;9(7):676–682. doi:10.1038/nmeth.2019

46. Taufalele PV, VanderBurgh JA, Muñoz A, Zanotelli MR, Reinhart-King CA. Fiber alignment drives changes in architectural and mechanical features in collagen matrices. PLoS One. 2019;14(5):e0216537. doi:10.1371/journal.pone.0216537

47. Tinevez JY, Perry N, Schindelin J, et al. TrackMate: An open and extensible platform for single-particle tracking. Methods. 2017;115:80–90. doi:10.1016/j.ymeth.2016.09.016

48. Comşa Ş, Cîmpean AM, Raica M. The Story of MCF-7 Breast Cancer Cell Line: 40 years of Experience in Research. Anticancer Res. 2015;35(6):3147–3154.

49. Chavez KJ, Garimella SV, Lipkowitz S. Triple negative breast cancer cell lines: One tool in the search for better treatment of triple negative breast cancer. Breast Disease. 2011;32(1-2):35–48. doi:10.3233/BD-2010-0307

50. Rubtsova SN, Zhitnyak IY, Gloushankova NA. Phenotypic Plasticity of Cancer Cells Based on Remodeling of the Actin Cytoskeleton and Adhesive Structures. International Journal of Molecular Sciences. 2021;22(4):1821. doi:10.3390/ijms22041821

51. Dongre A, Weinberg RA. New insights into the mechanisms of epithelial–mesenchymal transition and implications for cancer. Nat Rev Mol Cell Biol. 2019;20(2):69–84. doi:10.1038/s41580-018-0080-4

52. Lu W, Kang Y. Epithelial-Mesenchymal Plasticity in Cancer Progression and Metastasis. Developmental Cell. 2019;49(3):361–374. doi:10.1016/j.devcel.2019.04.010

53. Liu CY, Lin HH, Tang MJ, Wang YK. Vimentin contributes to epithelial-mesenchymal transition cancer cell mechanics by mediating cytoskeletal organization and focal adhesion maturation. Oncotarget. 2015;6(18):15966–15983. doi:10.18632/oncotarget.3862

54. Samani A, Zubovits J, Plewes D. Elastic moduli of normal and pathological human breast tissues: an inversion-technique-based investigation of 169 samples. Biomater Adv. 2007;52(6):1565–1576. doi:10.1088/0031-9155/52/6/002

55. Plodinec M, Loparic M, Monnier CA, et al. The nanomechanical signature of breast cancer. Nature Nanotech. 2012;7(11):757–765. doi:10.1038/nnano.2012.167

56. Desa DE, Wu W, Brown RM, et al. Second-Harmonic Generation Imaging Reveals Changes in Breast Tumor Collagen Induced by Neoadjuvant Chemotherapy. Cancers. 2022;14(4):857. doi:10.3390/cancers14040857

